# Ectoine production by newly isolated halophilic methanotroph *Methylotuvimicrobium percolatum* using surplus methane from anaerobic digestion

**DOI:** 10.1101/2025.09.13.675991

**Authors:** Shohei Yasuda, Ruri Ogasawara, Kazuma N. Fujii, Xinmin Zhan, Akihiko Terada

**Author notes:** Corresponding (Akihiko Terada).

## Abstract

Anaerobic digestion (AD) processes that convert waste such as livestock wastewater into biomethane for energy recovery are expanding globally. However, biogas exceeding storage capacity is typically flared, resulting in the release of carbon dioxide, a greenhouse gas. To address this issue, we focused on halophilic methane-oxidizing bacteria (methanotrophs) capable of converting methane into ectoine, a high-value compatible solute widely used in more than 400 cosmetic active ingredients and valued at approximately US$1,000/kg. Nevertheless, research on halophilic methanotrophs and ectoine biosynthesis remains limited. Here, we report the successful isolation of the Type I methanotroph *Methylotuvimicrobium percolatum* from activated sludge in a brackish water environment. Complete genome sequencing revealed genes for methane oxidation, osmoadaptation, and notably the *ectABCask* cluster responsible for ectoine biosynthesis. Activity tests under varying NaCl concentrations confirmed ectoine production, with a maximum yield of 20.1 mg ectoine/g-DCW achieved at 25% methane, 30 °C, and 3 wt% salinity. Biomass growth remained robust up to 6% salinity but declined sharply at higher NaCl concentrations, indicating that optimal conditions for growth and ectoine synthesis do not entirely coincide. These findings advance the fundamental understanding of halophilic methanotroph physiology and highlight their potential as sustainable platforms for converting surplus methane from AD processes into commercially valuable ectoine.

## Introduction

Anaerobic digestion (AD) processes, which extract biogas from waste such as livestock wastewater, are receiving growing attention, with global production expected to exceed 2,000 PJ/year by 2027 (International Energy Agency, 2023). Biogas contains 45–75% methane, which, when refined, is used for city gas and power generation (International Energy Agency, 2020). However, methane has a global warming potential 28 times higher than carbon dioxide and is the second most emitted anthropogenic greenhouse gas (IPCC, 2013). When biogas exceeds the storage capacity of AD facilities, it is flared, converting methane to carbon dioxide, a greenhouse gas, and releasing it into the atmosphere. In the United States alone, an estimated 1 billion cubic feet of recoverable methane from farms is combusted annually (U.S. Environmental Protection Agency, 2020). Together with possible incomplete incineration that causes methane emissions (Song et al., 2023), more effective methane utilization is required.

Halophilic microorganisms produce and accumulate compatible solutes as osmoprotectants that counteract the osmotic pressure in saline environments, and they adapt to environmental pressures by excreting various compatible solutes in response to increasing osmotic pressure (Czech et al., 2018). Compatible solutes include ectoine, hydroxyectoine, glycine betaine, proline, trehalose, and L-D-glutamate (Roberts, 2005). Among these, ectoine is commercially manufactured by the halophilic bacterium *Halomonas elongata* (Vandrich et al., 2020). Ectoine is also used as an active ingredient in many types of cosmetics and sunscreens. In contrast, commercial ectoine production remains costly because it requires high-quality glucose and oxygen, and equipment and substrates must be sterilized to prevent contamination (Pastor et al., 2010). Ectoine is of high commercial value, with a market price of approximately US$1,000/kg, and more efficient and cost-effective methods of bioproduction are required to reduce costs (Cantera et al., 2019; Strong et al., 2015).

Methanotrophs are gram-negative bacteria belonging to Proteobacteria, and they have attracted significant attention because they can assimilate methane as a carbon and energy source. Methanotrophs are classified as Type I or Type II based on their metabolic pathways. Type I methanotrophs mainly belong to Gammaproteobacteria and assimilate methane via the ribulose monophosphate methylerythritol phosphate (RuMP) pathway, while Type II methanotrophs belong to Alphaproteobacteria and use the serine pathway (Hanson and Hanson, 1996).

Some halophilic methanotrophs have been reported to produce valuable products, including ectoine. *Methylotuvimicrobium alcaliphilum* 20Z, a halophilic methanotroph, was reported to assimilate methane as a carbon source, subsequently synthesize and accumulate ectoine (Cantera et al., 2017; Khmelenina et al., 1999). Ectoine biosynthesis proceeds through transcription of the gene group *ectABCask*, which encodes the enzymes L-aspartokinase (Ask), L-2,4-diaminobutyric acid transaminase (EctB), L-2,4-diaminobutyric acid acetyltransferase (EctA), and L-ectoine synthase (EctC) (Reshetnikov et al., 2006). Halophilic methanotrophs may provide a means to utilize surplus biogas methane for ectoine production. However, research remains limited regarding other halophilic methanotrophs and ectoine production. Further research will provide fundamental knowledge to expand their practical applications.

This study aimed to isolate halophilic methanotrophs from a saline environment and conduct analyses to advance understanding of their physiology and ability to biosynthesize ectoine. Our previous shotgun metagenomic analysis revealed the ectoine biosynthesis genes in microbial communities inhabiting activated sludge treating landfill leachate [a brackish water environment with 1.23% salinity (Yasuda et al., 2021)], and we targeted halophilic methanotrophs for isolation from the microbiome. By obtaining the complete genome of this isolated methanotroph, we characterized its genetic potential for ectoine biosynthesis based on the presence of the *ectABCask* gene cluster and evaluated its ability to produce ectoine under varying salinity conditions. These findings provide fundamental insights into salt adaptation and ectoine biosynthesis in halophilic methanotrophs and support their potential application for upcycling surplus methane from anaerobic digestion processes.

## Materials and methods

### Isolation of methanotroph

Activated sludge treating brackish landfill leachate (Tokyo, Japan) (Yasuda et al., 2021) was collected. Methanotrophs were purified and isolated using a limiting dilution method. Activated sludge (20 μL) was mixed with DSMZ1180 mineral medium (180 μL) in microplate wells. DSMZ1180 medium was prepared by adding 30 g NaCl, 200 mg MgSO_4_·7H_2_O, 20 mg CaCl_2_·2H_2_O, 1 g KNO_3_, and 1 mL of trace element solution to 1 L of distilled water. The trace element solution was composed of 5 g EDTA, 100 mg CuCl_2_·5H_2_O, 2 g FeSO_4_·7H_2_O, 100 mg ZnSO_4_·7H_2_O, 20 mg NiCl_2_·6H_2_O, 200 mg CoCl_2_·6H_2_O, 30 mg Na_2_MoO_4_, 30 mg MnCl_2_·4H_2_O, and 30 mg H_3_BO_3_ in 1 L of distilled water. To maintain the pH at 8.5–9.0, carbonate buffer (50 mL/L 1 M NaHCO_3_; 5 mL/L 1 M Na_2_CO_3_) and phosphate buffer (14 g KH_2_PO_4_, 30 g Na_2_HPO_4_·12H_2_O, 1 L distilled water) were added at 20 mL/L. The culture medium was serially diluted 10-fold in a microplate and cultured in an AnaeroPack™ (Mitsubishi Gas Chemical Company, Inc., Tokyo, Japan) containing 20% methane and 80% air. The most diluted cell cultures that showed turbidity were transferred to a new microplate, and the same procedure was repeated. To periodically verify the purity of the isolate, DNA was extracted using the FastDNA Spin Kit for Soil (MPBiomedicals, Santa Ana, CA, USA) and the V4 region of the 16S rRNA gene was amplified with universal primers (515f and 806r^15^). Sanger sequencing was performed using a 3730xl DNA Analyzer (ThermoFisher Scientific Inc., MS, US) at an analysis service (Fasmac Co., Ltd., Kanagawa, Japan), and the purity was confirmed by examining chromatogram waveforms. Finally, isolation was verified by analyzing the banding pattern using polymerase chain reaction–restriction fragment length polymorphism (PCR-RFLP).

### Preculture of isolated methanotroph

Preculturing of the isolate was conducted by placing 30 mL DSMZ1180 medium (3% NaCl) in a 120 mL vial sealed with a butyl rubber stopper and aluminum cap, and 30 mL of methane was injected via a syringe. The isolated strain was added using a syringe and cultured at 30 °C and 130 rpm shaking for 10 days. All operations were conducted under sterile conditions.

### Morphological observation of the isolated methanotroph by scanning electron microscopy

Cell morphology and attachment of the isolated strain were examined by scanning electron microscopy (SEM) using sponge carriers (1 cm³) grafted with diethylamino functional groups to enhance microbial adhesion (Terada et al., 2012). Cells grown in DSMZ1180 medium (3% NaCl) for 5 days were incubated with the carriers (30 mL, 24 h) and fixed with 4% paraformaldehyde containing 25% glutaraldehyde at 4 °C for 2 h. After three washes in PBS/CaCl₂ solution, samples were dehydrated through a graded ethanol series (25–100%) and dried at 46 °C. The dried carriers were split, mounted on SEM stubs with conductive tape, sputter-coated with gold (1 min; Quick Coater SC-701 MKⅡ ECO, SANYU ELECTRON), and observed with a Miniscope TM3030 (HITACHI, Tokyo, Japan).

### DNA extraction, sequencing, and hybrid assembly

Genome analysis was performed using the precultured biomass of the isolate. The genomic DNA of the isolate was extracted with a phenol-chloroform extraction and RNA was removed using RNaseA (TaKaRa Bio, Inc., Japan). Long reads were not fragmented, and libraries were prepared using a 1D ligation sequencing kit (SQK-LSK-109; Oxford Nanopore Technologies Ltd., Oxford, UK). Sequencing was performed using a MinION Mk1B (FLO-MIN110; Oxford Nanopore Technologies Ltd., Oxford, UK) with an R.10 flow cell, and sequence data were base-called with Guppy v5.0.11 (https://nanoporetech.com/document/Guppy-protocol). Sequence quality was checked with NanoPlot v1.40.0 (De Coster and Rademakers, 2023), and adapter sequences, low-quality reads (<Q7), headers (75 bp), and short reads (<30,000 bp) were removed with NanoFilt v2.8.0 (De Coster and Rademakers, 2023). For short-read sequencing, library preparation was performed according to the manufacturer’s protocol using MGI Easy FS PCR Free DNA Library Prep Set (MGI Tech., Shenzhen, China). Libraries were created by a sequencing service (Genome-Lead Co., Ltd., Kagawa, Japan) and 150 bp paired-end sequencing was performed using DNBSEQ-G400FAST (MGI Tech., Shenzhen, China). Adapter sequences, barcode sequences, low-quality reads (<Q30), 1 base pair at the end, reads <20 bp, and reads with N ≥ 10 were removed using Fastp v0.23.2 (Chen et al., 2018). Hybrid assembly was performed by Unicycler v0.5.0 (Wick et al., 2017) using long and short reads. The completeness of the obtained genome was evaluated using gVolante v2.0.0 (Nishimura et al., 2017) with the BUSCO v5 pipeline (Manni et al., 2021). Clusters of orthologous genes (COGs), coding sequence (CDS), tRNA, rRNA, GC content, and GC skew of the obtained genome were visualized using Genovi v0.4.3 (Cumsille et al., 2023).

### Phylogenetic analysis and genome annotation

Phylogenetic analysis was performed by GTDB-Tk v2.4.1 (Chaumeil et al., 2020) using classify_wf (reference data version r226), and a phylogenetic tree was created for the family *Methylomonadaceae* using de_novo_wf with 120 marker genes (outgroup_taxon: g Methylotetracoccus; taxa_filter: f Methylomonadaceae). To confirm the novelty of the isolate, the average nucleotide identity (ANI) with the closest related species was calculated using skani v0.2.2 (Shaw and Yu, 2023) included in classify_wf. Genome annotation was performed using DFAST v1.2.17 (Tanizawa et al., 2018) (--fix_origin --offset 100 --aligner blastp). The obtained amino acid sequences were annotated by mapping them to the reference metabolic pathway of the Kyoto Encyclopedia of Genes and Genomes (KEGG; release 2023-04-01) (Kanehisa and Goto, 2000) using BlastKOALA v3.0 (Kanehisa et al., 2016), and the functional genes in the isolate were identified. Because the isolate was expected to be a methanotroph and inhabit a saline environment, genes encoding methane oxidation and compatible solute biosynthesis (e.g., ectoine, hydroxyectoine, glycine betaine, proline, trehalose, and L-D-glutamate) were investigated, along with genes involved in central metabolism, including the tricarboxylic acid (TCA) cycle, glycolysis, carbon, and nitrogen metabolism. Default parameters were used for all software unless otherwise specified.

### Activity tests of the isolated methanotroph under varying salinity

To evaluate the effects of salinity on growth and ectoine production, activity tests were conducted by varying NaCl concentration (3–12%) while maintaining constant methane and oxygen conditions (25% methane balanced with air, corresponding to ∼15–16% O₂) at 30 °C. Culture vials were prepared following the same procedure as the preculture, except for the adjusted salinity. Each vial was aseptically inoculated with 100 μL of the precultured strain (OD₅₉₀ = 0.08). All experiments were performed in triplicate, and headspace gas and culture medium were periodically sampled to determine dry cell weight (DCW), dissolved gas concentrations, and ectoine content.

### Measurement of dissolved gas concentration in the medium

During the activity test, we analyzed the methane oxidation ability of the isolate. The concentrations of dissolved methane, oxygen, and carbon dioxide in the medium were calculated by measuring the respective gas concentrations in the headspace gas and converting them to dissolved concentrations using Henry’s constants. The gas concentrations in the headspace of each vial were measured by gas chromatography–mass spectrometry (GC-MS-QP2010 Plus, Shimadzu, Kyoto, Japan) using a CP-PoraPLOT column (film thickness 10 µm, length 25 cm, and inner diameter 320 µm, Agilent Technologies, Tokyo, Japan). After measuring the internal pressure of the vial using a manometer (PG-100-101GP, Nidec Copal Electronics, Tokyo, Japan), 200 μL of headspace gas was collected from the vial using a syringe and directly injected into the GC-MS. Helium was used as the carrier gas at a flow rate of 25.3 mL/min. The temperature of the inlet port, column, ionization source of the mass detector, and interface were set at 100 °C, 50 °C, 200 °C, and 250 °C, respectively. The concentrations of methane, oxygen, and carbon dioxide in the headspace were measured with reference to their respective standard gases (G1 grade, Taiyo Nippon Sanso Corporation, Tokyo, Japan), and the partial pressures of each gas were calculated from the measured internal pressure of the vials to obtain the molar fractions in the headspace. The molar fractions of gases in the medium were calculated using Henry’s constants. Henry’s constants in high saline conditions were calculated from the Sechenov equation (Eq. 1), which correlates the values of nonpolar gases, such as methane, to the salinity. In all calculations, the salt concentration, the decrease in the liquid volume resulting from liquid sample collection, and the increase in the headspace volume were considered. The methane oxidation rate was calculated from the change in methane concentration during the logarithmic growth phase.

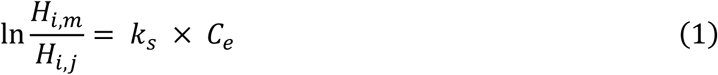

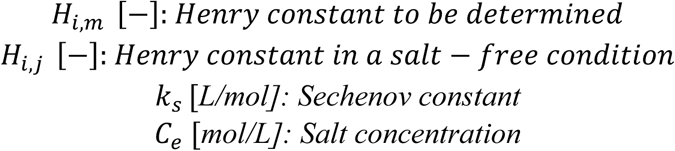

### Dry cell weight measurement

The optical density (OD_590_) value of the medium was converted to the DCW of the isolate in the activity test. The biomass from the logarithmic growth phase onwards was serially diluted with DSMZ1180 medium in preculture to attain the conversion factor. After measuring the OD_590_, the cells were freeze dried and weighed to obtain the correlation between OD_590_ and cell weight. In the activity test, 2 mL of the culture solution was sampled over time, followed by OD_590_ measurement and DCW conversion. The maximum growth rate was calculated from the change in the DCW during the logarithmic growth phase of the isolate. The maximum specific growth rate was calculated by dividing the maximum growth rate by the DCW.

### Ectoine quantification by HPLC

After OD measurement, 2 mL of the culture solution was centrifuged at 9,000 rpm and 4 °C for 15 min, and the resulting pellet was resuspended in 1 mL of 80% ethanol and transferred to a bead milling tube. The sample was centrifuged again at 10,000 rpm and 4 °C for 15 min, the supernatant was discarded, and 1 mL 80% ethanol was added to resuspend the culture solution. The mixture was homogenized with a FastPrep instrument (FastPrep 24 Instrument Version 4, MP Biomedicals, Japan K. K., Tokyo, Japan) at 6 m/s for 60 seconds and incubated at room temperature overnight. Then, the mixture was centrifuged at 10,000 rpm and 4 °C for 15 min. The supernatant was filtered through a 220-nm filter and refrigerated until HPLC measurement. Before HPLC analysis, the sample was diluted 10-fold with 70% acetonitrile aqueous solution. An ectoine standard (> 98.0% purity; Tokyo Chemical Industry Co., Ltd., Tokyo, Japan) at 10,000 mg/L was serially diluted with 70% acetonitrile for calibration. Ectoine was measured by HPLC with a UV detector (Shimadzu, Kyoto, Japan) and C18 column (Supelco™ Ascentis™, 5 μm, 15 cm × 4.6 mm, Merck Electronics Ltd., Tokyo, Japan). The column temperature was 40 °C, and the eluent, consisting of acetonitrile and 0.1% v/v phosphoric acid aqueous solution, was supplied at a flow rate of 1 mL/min. From the start of the measurement, acetonitrile and 0.1% v/v phosphoric acid aqueous solution were run at a ratio of 1:99 for the first 5 minutes, followed by a gradient mode in which the ratio increased to 95:5 in 20 minutes. A 20 μL portion of the sample was injected and measured by HPLC. All analyses were visualized by R v4.4.2 (R Core Team, 2024) using ggplot2 v2.3.5.1 (Wickham, 2016). The ectoine yield was measured using the ectoine quantity in DCW (Eq. 2).

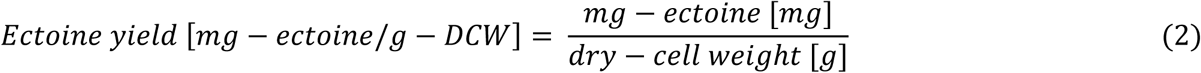

## Results and discussion

### Isolated strains and genomic analysis

This study reports the first successful isolation of an ectoine-producing halophilic methanotroph, *Methylotuvimicrobium percolatum*, from a brackish environment. Isolation of a methanotroph is a crucial step toward utilizing surplus methane from AD plants for the biosynthesis of value-added products. A single microorganism was obtained after eight successive rounds of cultivation and transfer using the limiting dilution method applied to activated sludge from a brackish water environment. SEM observations revealed that the isolated cells were attached to the carrier surface and exhibited an oval shape, measuring approximately 1.25–1.8 µm in length and 0.6–0.75 µm in width, with smooth cell surfaces (**Fig. 1a**). Genomic DNA extracted from the isolate was sequenced using long- and short-read platforms and subjected to hybrid assembly. After removal of adapters and low-quality reads, 35.6 K long reads (1.6 Gbp) and 8.8 M short reads (1.3 Gbp) were assembled into a circular chromosome (5.3 Mbp) and a circular plasmid (119.9 Kbp) (**Fig. 1b**). The assembled genome had a GC content of 48.6%, encoded 4,512 coding sequences (CDSs), nine rRNAs, and 44 tRNAs, and exhibited 100% completeness. Phylogenetic analysis placed the isolate within the genus *Methylotuvimicrobium* (family *Methylomonadaceae*), a Type I methanotroph most closely related to *Methylotuvimicrobium* sp. 018830325 (Gammaproteobacteria bacterium; NCBI taxonomy) (**Fig. 2**). The average nucleotide identity (ANI) with strain 018830325 was 98.3%, exceeding the 95% threshold for species delineation, confirming that the isolate belongs to the same species. We propose the name *Methylotuvimicrobium percolatum* (per’co.la.tam., referring to “leachate”).

**Fig. 1.**
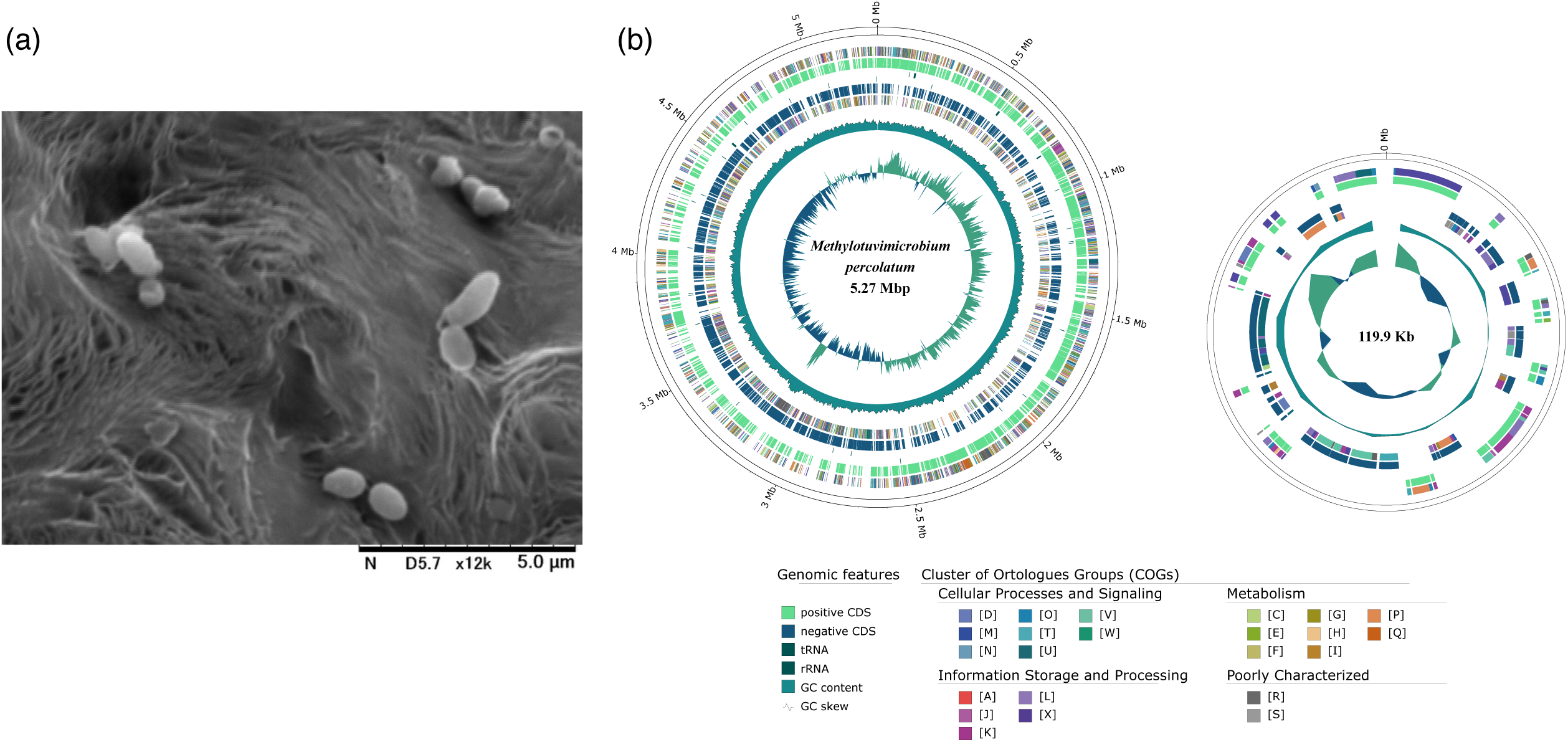
Cell morphology and genome organization of isolated *Methylotuvimicrobium percolatum*. (a) Scanning electron micrograph showing oval-shaped cells (length 1.25–1.8 µm, width 0.6–0.75 µm) attached to the carrier surface. (b) Circular genome (5.27 Mbp) and plasmid (119.9 Kbp) organization, illustrating clusters of orthologous genes (COGs), coding sequences (CDSs), tRNAs, rRNAs, GC content, and GC skew.

**Fig. 2.**
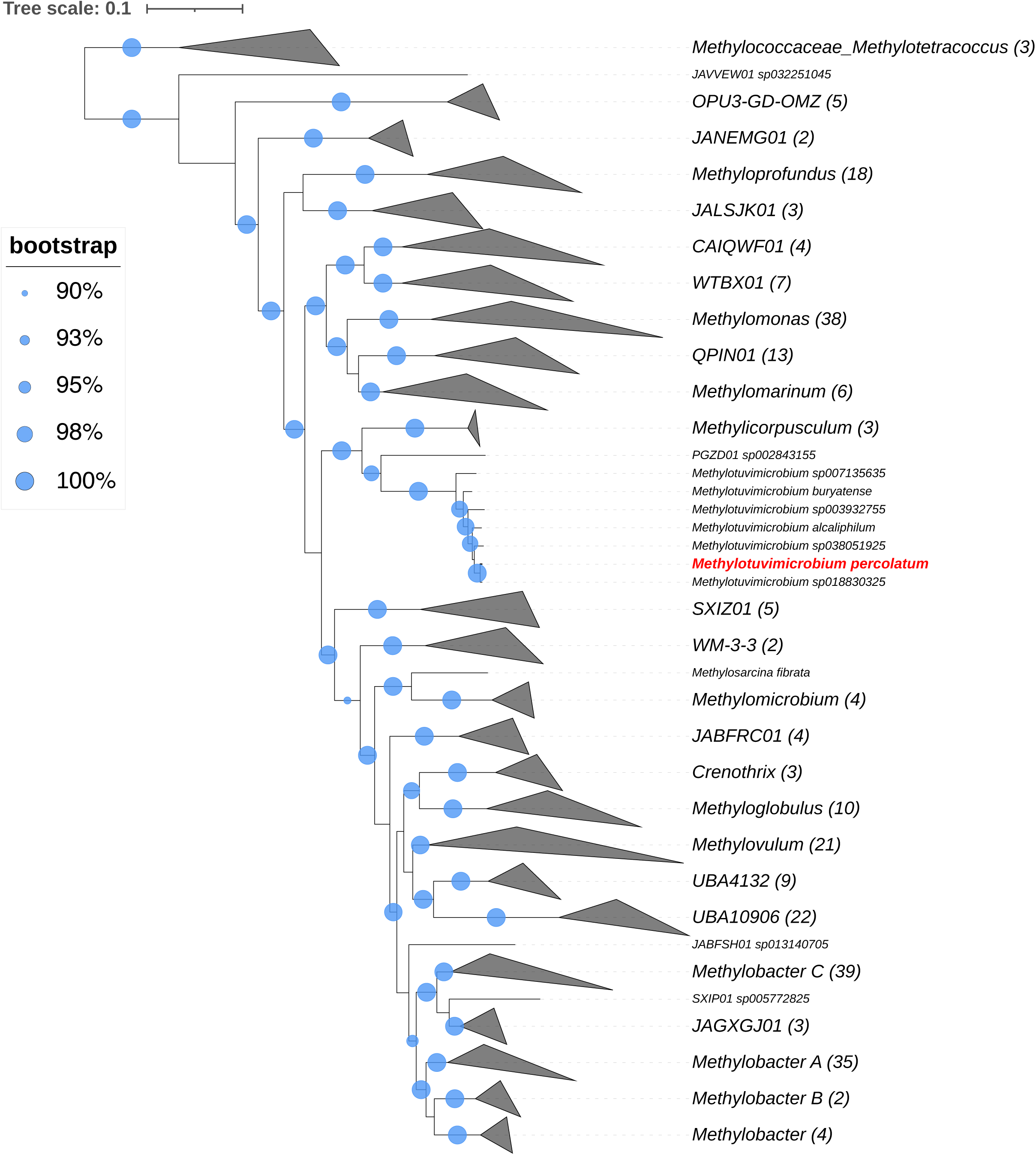
Phylogenomic placement of the *M. percolatum* genome within the family *Methylomonadaceae*, inferred from the concatenated alignment of 120 marker genes. The tree is rooted with *Methylotetracoccus*. The *M. percolatum* genome is highlighted in red. Bootstrap support values of 90–100% are denoted by light blue circles. Numbers in parentheses indicate leaves within the collapsed nodes.

### Genetic function analysis of *M. percolatum*

The predicted amino acid sequences of *M. percolatum* were annotated against KEGG reference pathways to clarify its metabolic potential (**Table 1**). The genome encodes the *pmoABC* gene cluster for particulate methane monooxygenase, *mxaACDFGIJKL* and *xoxF* for methanol dehydrogenase, and *fdhF*, *fdsD*, and *fdwB* for formate dehydrogenase, indicating a complete pathway for methane oxidation to CO_2_. Genes linking formaldehyde metabolism through the RuMP pathway to the tricarboxylic acid (TCA) cycle and glycolysis were also present, corroborating the classification of *M. percolatum* as a Type I methanotroph suggested by the phylogenomic analysis. The genome also contains the *ectABCask* gene cluster responsible for ectoine biosynthesis, confirming its genetic potential to produce ectoine. Notably, genes associated with ectoine biodegradation (*doeAB*) were absent, suggesting that synthesized ectoine is likely retained intracellularly. In addition, genes for biosynthesis of other compatible solutes—including *ectD* (hydroxyectoine), *proABC* and *putA* (proline), *treSYZ* (trehalose), and *GDH2* and *gltB* (L-D-glutamate) —were identified, whereas genes for glycine betaine biosynthesis were not detected. The genome harbors multiple ion transporter genes that support osmoadaptation, including *clcA* (chloride ion channel), *nqrABCDE* (sodium-translocating NADH:quinone oxidoreductase), *trkH, ktrA,* and *trkA* (potassium transporters), and *aqpZ* (aquaporin). These genes likely contribute to maintaining osmotic balance in high-salinity environments. The presence of both ectoine biosynthesis genes and diverse ion transport systems suggests that *M. percolatum* is well adapted to fluctuating saline conditions and may be suitable for biotechnological applications involving saline methane streams.

**Table 1.**
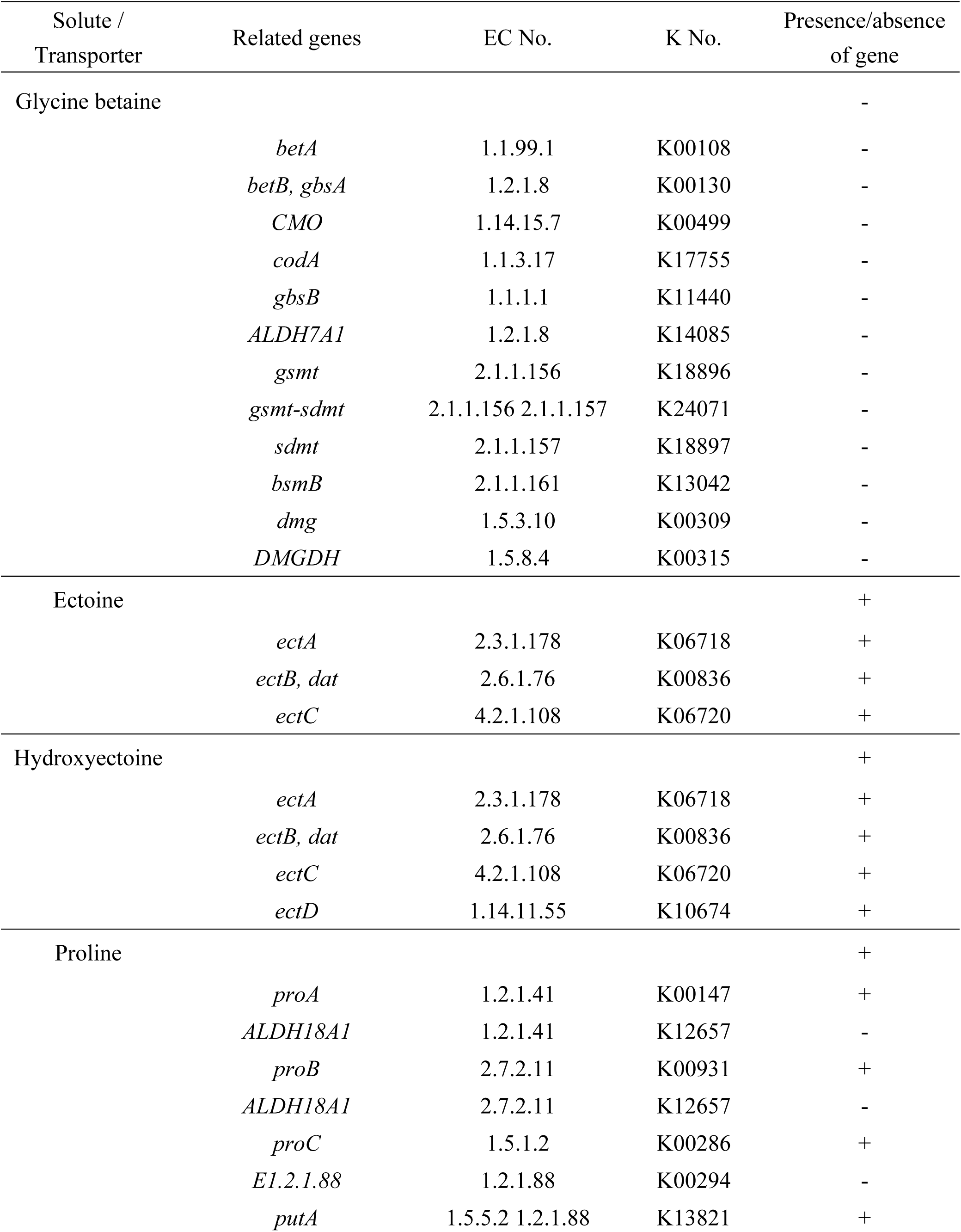
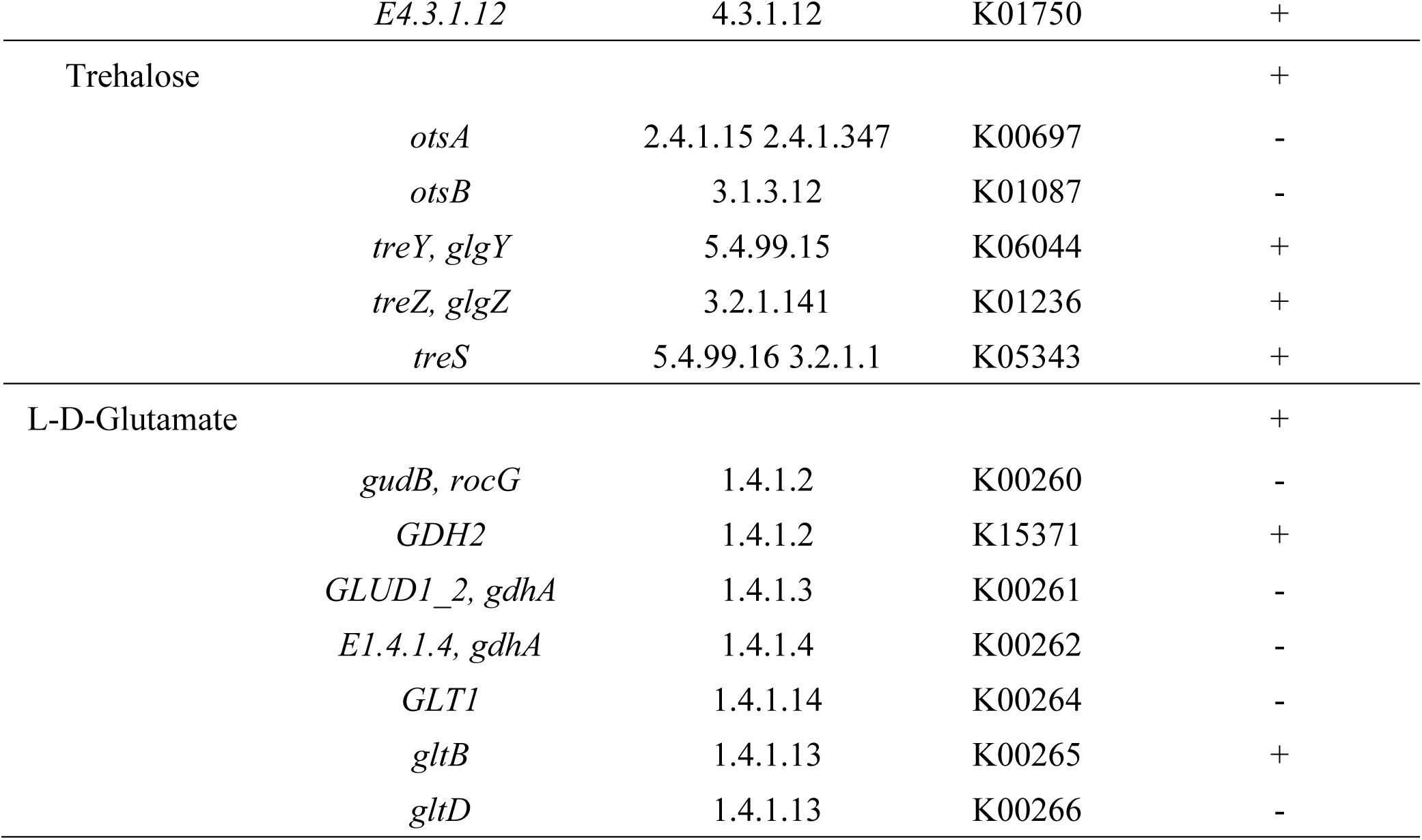
Presence and absence of genes related to osmolyte biosynthesis in *M. percolatum*. Enzyme Commission (EC) numbers and KEGG Orthology (K) numbers are indicated.

The type strain of the genus *Methylotuvimicrobium* is *M. alcaliphilum* 20Z, and research on ectoine biosynthesis using methane as a substrate has recently emerged (Reshetnikov et al., 2006; Cantera et al., 2019, 2017). However, only five species of this genus have been described to date (Kaluzhnaya et al., 2001; Kalyuzhnaya et al., 2008; Orata et al., 2018), and only four of them have fully sequenced genomes (Groom et al., 2019; Khmelenina et al., 2013; Vuilleumier et al., 2012). Thus, the isolation of *M. percolatum*, together with the analysis of its physiological traits, particularly methane oxidation and ectoine production, and the determination of its genetic potential for biosynthesis based on the complete genome, represents a significant contribution to advancing knowledge of ectoine production by methanotrophs.

### Activity test for ectoine biosynthesis and changes in ectoine yield over time

Ectoine production and biomass accumulation by *Methylotuvimicrobium percolatum* were evaluated under varying NaCl concentrations (3–12%) while maintaining constant methane and oxygen conditions (25% methane balanced with air, corresponding to ∼15–16% O₂) at 30 °C (**Fig. 3**). Maximum DCW reached 132 ± 3 mg/L at 3% NaCl and remained comparable at 6% NaCl (129 ± 5 mg/L), but sharply decreased to ∼14 mg/L at 9% and 12% NaCl. Specific growth rate followed a similar trend, peaking at 0.42 ± 0.06 day⁻¹ at 6% NaCl and dropping markedly at higher salinities.

**Fig. 3.**
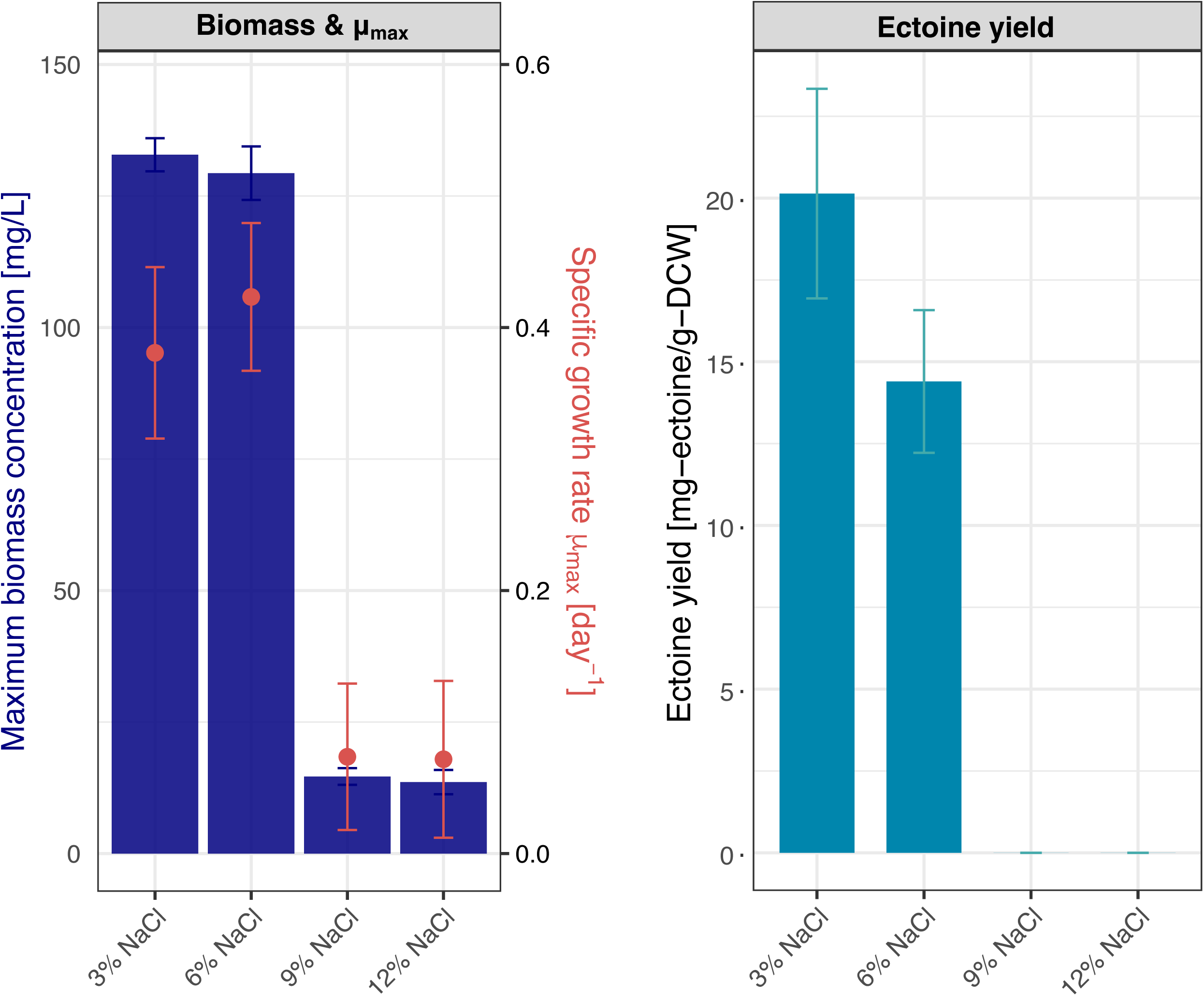
Maximum biomass concentration (DCW), specific growth rate (*μ*_max_), and ectoine yield of *M. percolatum* under varying NaCl concentrations (3–12%) at 25% methane and 30 °C.

Ectoine yield exhibited a distinct response to salinity. The highest yield was observed at 3% NaCl (20.1 ± 3.2 mg-ectoine/g-DCW), decreased to 14.4 ± 2.2 mg/g-DCW at 6%, and was undetectable at ≥ 9%. These results indicate that moderate salinity (3–6%) enhances ectoine biosynthesis, whereas excessive salinity inhibits both growth and compatible solute production. Although the maximum yield obtained here was lower than that reported for *Halomonas elongata* (155 mg/g-DCW at 15% salt) (Sauer and Galinski, 1998), further optimization—such as adjusting salinity, supplementing trace elements like tungsten or copper known to enhance methanotroph growth, and refining temperature conditions—may improve production. Previous studies report ectoine synthase activity peaking at 15 °C (Czech et al., 2018; Widderich et al., 2016), while others observed higher yields at 25 °C (Cantera et al., 2016), highlighting the need to examine the interplay between temperature and salinity for maximizing ectoine production.

In commercial production, the salt concentration is typically set at a high level and subsequently replaced with a low-salt medium (preferably salt-free) to reduce the osmotic pressure. The “bacterial milking” method is used to release the ectoine accumulated intracellularly in *H. elongata* cells (Sauer and Galinski, 1998). Destroying bacterial cells to extract intracellularly accumulated compounds (*e.g.*, ectoine) requires the entire process of cultivation to be repeated for continuous production, but bacterial milking avoids cell disruption and recultivation and retains bacterial physiological activity by extracting ectoine using changing salt concentrations in the culture medium. *M. percolatum* has a group of genes encoding chloride, sodium, and potassium transporters, as well as aquaporin, suggesting that it has the capability to adjust to the osmotic pressure it is exposed to. Therefore, *M. percolatum* could possibly withstand the fluctuating salinity conditions during the bacterial milking process. Future studies are required to determine the optimal environment to efficiently and sustainably produce ectoine using *M. percolatum*.

## Conclusion

In this study, the ectoine-producing methanotroph, *Methylotuvimicrobium percolatum*, was successfully isolated from a brackish landfill leachate environment, and its genome and physiological traits were comprehensively characterized. Genome analysis revealed the presence of the *ectABCask* gene cluster, indicating the genetic potential for ectoine biosynthesis. Cultivation tests conducted at 30 °C and 25% methane while varying NaCl concentrations (3–12%) demonstrated that biomass (132 mg DCW/L) and ectoine yield (20.1 mg/g-DCW) peaked at 3% NaCl, whereas growth and production were strongly inhibited at ≥9% NaCl. These findings indicate that moderate salinity favors ectoine synthesis, while optimal conditions for growth and solute production do not necessarily coincide. This work provides fundamental insights into methane-based production of compatible solutes, highlighting a new biotechnological route for upcycling surplus methane into high-value ectoine. Future studies should optimize cultivation parameters, including salinity, temperature, and trace elements, and evaluate the applicability of extraction techniques such as bacterial milking to establish a sustainable and scalable ectoine production process.

## CRediT authorship contribution statement

**Shohei Yasuda:** Writing – original draft, Conceptualization, Resources, Methodology, Software, Validation, Formal analysis, Investigation, Data curation, Visualization, Funding acquisition. **Ruri Ogasawara:** Methodology, Validation, Formal analysis, Investigation, Data curation. **Kazuma N. Fujii:** Methodology, Validation, Formal analysis. **Xinmin Zhan:** Writing – review & editing. **Akihiko Terada:** Writing – review & editing, Conceptualization, Methodology, Resources, Supervision, Project administration, Funding acquisition.

## Acknowledgments

We thank the late Ms. Kanako Mori for her experimental support, Dr. Kuninori Morimoto for his computational analysis support, Prof. Hideaki Tokuyama for cell observation by scanning electron microscopy, and Dr. Kousho Hosoi for fruitful discussion. This work was supported by KAKENHI Grant-in-Aid for Scientific Research (22K14348, 23H03565, and 23K28255) from Japan Society for the Promotion of Science, Japan Science and Technology Agency grant number JPMJPF2104, and Kurita Water and Environment Foundation (21A043 and 22T012).

## Data availability

The complete genome sequence of *Methylotuvimicrobium percolatum* has been deposited in DDBJ/EMBL/GenBank under BioProject accession number PRJDB19528 and Run accession number DRR623457-DRR623458.

